# BERTology Meets Biology: Interpreting Attention in Protein Language Models

**DOI:** 10.1101/2020.06.26.174417

**Authors:** Jesse Vig, Ali Madani, Lav R. Varshney, Caiming Xiong, Richard Socher, Nazneen Fatema Rajani

## Abstract

Transformer architectures have proven to learn useful representations for protein classification and generation tasks. However, these representations present challenges in interpretability. Through the lens of attention, we analyze the inner workings of the Transformer and explore how the model discerns structural and functional properties of proteins. We show that attention (1) captures the folding structure of proteins, connecting amino acids that are far apart in the underlying sequence, but spatially close in the three-dimensional structure, (2) targets binding sites, a key functional component of proteins, and (3) focuses on progressively more complex biophysical properties with increasing layer depth. We also present a three-dimensional visualization of the interaction between attention and protein structure. Our findings align with known biological processes and provide a tool to aid discovery in protein engineering and synthetic biology. The code for visualization and analysis is available at https://github.com/salesforce/provis.

## 1 Introduction

The study of proteins, the fundamental macromolecules governing biology and life itself, has led to remarkable advances in understanding human health and the development of disease therapies. Protein science, and especially protein engineering, has historically been driven by experimental, wetlab methodologies along with biophysical, structure-based intuitions and computational techniques [6, 33, 69]. The decreasing cost of sequencing technology has enabled us to collect vast databases of naturally occurring proteins [21], which are rich in information that is ripe for developing powerful sequence-based approaches. For example, sequence models leveraging principles of co-evolution, whether modeling pairwise or higher-order interactions, have enabled prediction of structure or function [66].

Proteins, as a sequence of amino acids, can be viewed precisely as a language. As such, they may be modeled using neural architectures that have been developed for natural language. In particular, the Transformer [80], which has led to a revolution in unsupervised learning for text, shows promise for a similar impact on protein sequence modeling. However, the strong performance of the Transformer comes at the cost of interpretability, and this lack of transparency can hide underlying problems such as model bias and spurious correlations [41, 55, 75]. In response, a great deal of NLP research now focuses on interpreting the Transformer, e.g., the subspecialty of “BERTology” [65], which studies the BERT [19] Transformer model specifically.

In this work, we adapt and extend this line of interpretability research to protein sequences. We analyze a Transformer protein model through the lens of attention, the core component of the Transformer architecture, and we present a set of analysis methods that capture the unique functional and structural characteristics of protein sequences. In contrast to NLP, which aims to automate a capability that humans already possess—understanding natural language—protein modeling also seeks to shed light on biological processes that are not yet fully understood. Therefore we also discuss how interpretability can aid scientific discovery.

Our analysis reveals that attention captures high-level structural properties of proteins, connecting amino acids that are spatially close in three-dimensional structure, but far apart in the underlying sequence (Figure 1a). We also find that attention targets binding sites, a key functional component of proteins (Figure 1b). Further, we show that attention captures substitution properties of amino acids, and constructs progressively higher-level representations of structure and function with increasing layer depth.

**Figure 1:**
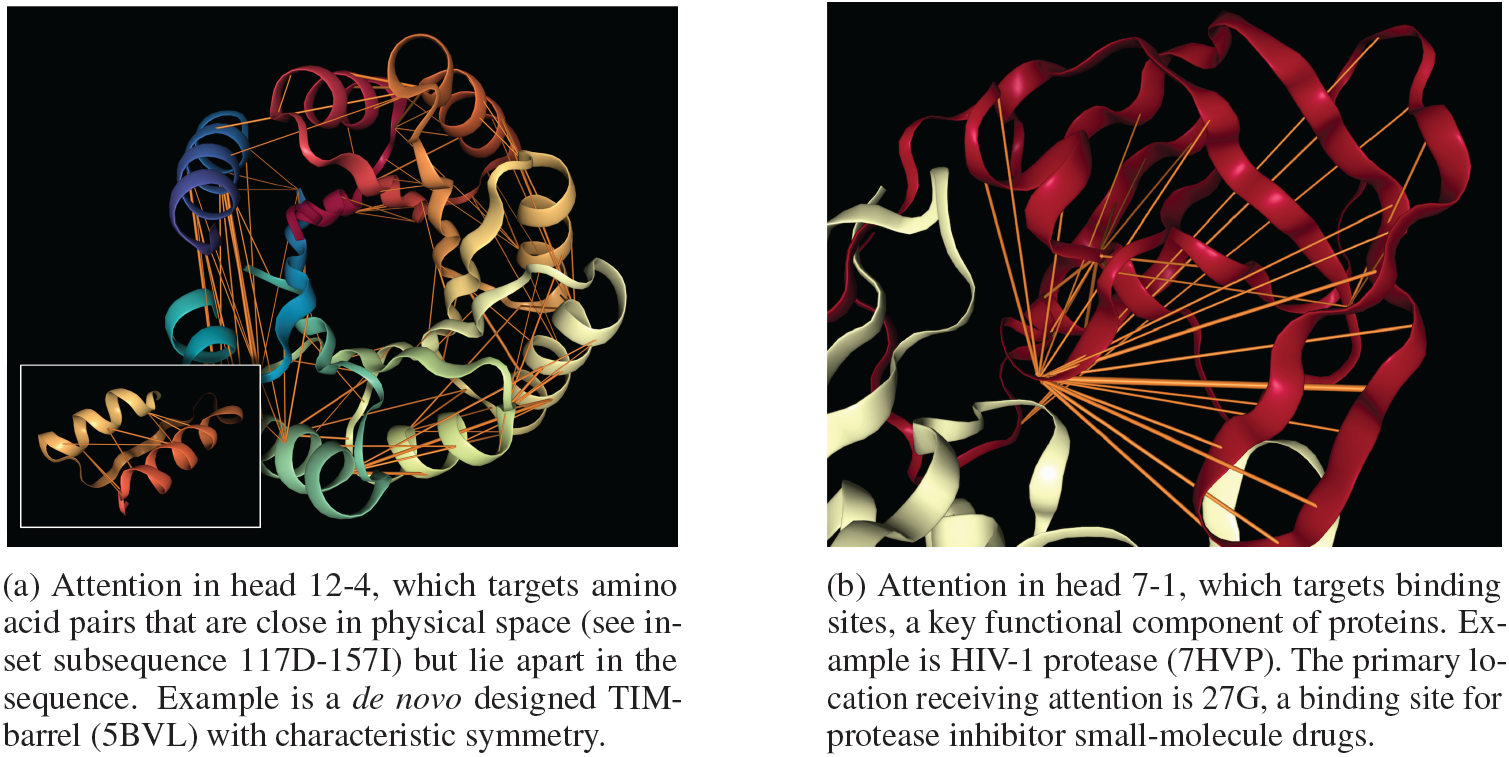
Two examples of how specialized attention heads in a Transformer recover protein structure and function, based solely on language model pre-training. The orange lines depict attention between amino acids, with line width proportional to attention weight. Heads were selected based on alignment with ground-truth annotations of contact maps and binding sites. Attention values below 0.1 are hidden. Visualizations based on the NGL Viewer [54, 67, 68].

## 2. Background: Proteins

In this section we provide background on the biological concepts discussed in later sections.

### Amino acids

Protein sequences are composed from a vocabulary of 20 standard amino acids. Just as different words may convey similar meanings, different amino acids may play similar structural and functional roles in a sequence. While word synonyms are encoded in a thesaurus, similar amino acids are captured in a *substitution matrix*, which scores pairs of amino acids on how readily they may be substituted for one another while maintaining protein viability. One common substitution matrix is BLOSUM [29], which is derived from co-occurrence statistics of amino acids in aligned protein sequences.

### Secondary and tertiary structure

Though a protein may be abstracted as a sequence of amino acids, it represents a physical entity with well-defined three-dimensional structure (Figure 1). *Secondary structure* describes the local segments of proteins, and may be divided into three broad categories: *Helix, Strand*, and *Turn/Fold*, as well as an *Other* category for local structure that does not fall into one of these categories. *Tertiary structure* encompasses the large-scale formations that determine the overall shape and function of the protein. One way to characterize tertiary structure is by a *contact map*, which describes the pairs of amino acids that are in contact (within 8 angstroms of one another) in the folded protein structure but lie apart (by at least 6 positions) in the underlying sequence [60].

### Binding sites

Proteins may also be characterized by their functional properties. *Binding sites* are regions of protein sequences that bind with other molecules to carry out a specific function. For example, the HIV-1 protease is an enzyme responsible for a critical process in replication of HIV [11]. It has a binding site shown in Figure 1b which is a target for drug development to ensure inhibition.

## 3. Methodology

### Model

We study the BERT Transformer model [19], though this analysis could be applied to other Transformers as well. In particular, we use the BERT-Base model from the TAPE repository [60], which was pretrained on masked language modeling of amino acids over a dataset of 31 million protein sequences [22]. This model accepts as input a sequence of amino acids ***x*** = (*x*_1_, …, *x*_*L*_) and outputs a sequence of continuous embeddings ***z*** = (*z*_1_, …, *z*_*L*_). The architecture comprises a series of encoder layers, each of which includes multiple attention heads. Each attention head generates a distinct set of attention weights *α* for an input, where *α*_*i,j*_ > 0 represents the attention from token *i* to token *j* in the sequence, such that *j α*_*i,j*_ = 1. Intuitively, attention weights define the influence of every token on the next layer’s representation for the current token. The BERT-Base model has 12 layers and 12 heads, yielding a total of 144 distinct attention mechanism. We denote a particular layer-head pair by *<layer>-<head_index>*, e.g. head *3-7* for the 3rd layer’s 7th attention head. A detailed formulation of the attention mechanism may be found in Appendix A.1.

### Attention analysis

We analyze how attention aligns with various protein properties, both at the token level (secondary structure, binding sites) and at the token-pair level (contact maps). In the latter case, we define an indicator function *f* (*i, j*) for property *f* that returns 1 if the property is present in token pair (*i, j*) (i.e., if amino acids *i* and *j* are in contact), and zero otherwise. We then compute the proportion of attention that aligns with *f* over a dataset ***X*** as follows:

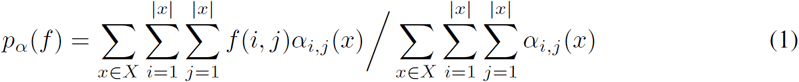

where *α*_*i,j*_(*x*) denotes the attention from *i* to *j* for input sequence *x*.

For a *token*-level property *f*, we define *f* (*i, j*) to be an indicator that returns 1 if the property is present in token *j* (e.g. if *j* is a binding site) and zero otherwise. In this case, *p*_*α*_(*f*) represents the proportion of attention that is directed *to* property *f*.

### Probing tasks

Besides analyzing attention, we also use a diagnostic classifier [1, 16, 81] to probe the layer outputs to determine what information they contain about the properties described above. For the classifier, we use a single linear layer followed by softmax, and we freeze the weights of the original model. For token-level probing tasks (binding sites, secondary structure), we feed each token’s output vector directly to the classifier. For token-pair probing tasks (contact map), we construct a pairwise feature vector by concatenating the elementwise differences and products of the two tokens’ output vectors, following the TAPE^1^ implementation. We use task-specific evaluation metrics: for secondary structure prediction, we measure F1 score; for contact prediction; we measure precision@*L/*5, where *L* is the length of the protein sequence, following standard practice [52]; for binding site prediction, we measure precision@*L/*20, since approximately one in twenty amino acids in each sequence is a binding site (4.8% in the dataset described below).

### Datasets

We use two protein sequence datasets from the TAPE repository for the analysis: the ProteinNet dataset [3, 10, 26, 52] and the Secondary Structure dataset [10, 38, 52, 60]. Both datasets contain amino acid sequences. ProteinNet is also annotated with spatial coordinates of each amino acid, which is used for generating contact maps, and the Secondary Structure dataset is annotated with the secondary structure at each sequence position. For our analyses of amino acids and contact maps, we use ProteinNet, and for the analysis of secondary structure and binding sites we use the Secondary Structure dataset. For the binding site analysis, we also obtained available token-level binding site annotations from the Protein Data Bank [10]. For analyzing attention, we used a random subset of 5000 sequences from the respective training splits as this analysis was purely evaluative; for training the probing classifier, we used the full training splits for training the model, and the validation splits for evaluation. More details about the datasets may be found in Appendix A.2.

### Experimental details

We exclude attention to the [SEP] delimiter token, as it has been shown to be a “no-op” attention token [15], as well as attention to the [CLS] token, which is not explicitly used in language modeling. We also filter attention below a minimum threshold of 0.1 (on a scale from 0 to 1) to reduce the effects of very low-confidence attention patterns on the analysis. We truncate all protein sequences to a maximum length of 512 to reduce the model memory requirements.2 Experiments are performed using a single Tesla V-100 GPU with 16GB memory.

## 4. What does attention understand about proteins?

### 4.1 Amino Acids

We begin by examining the interaction between attention and particular types of amino acids.

#### Attention heads specialize in certain types of amino acids

We computed the proportion of attention that each head focuses on particular types of amino acids, averaged over a dataset of 5000 sequences with a combined length of 1,067,712 amino acids. We found that for 14 of the 20 types of amino acids, there exists an attention head that focuses over 25% of attention on that amino acid. For example, Figure 2 shows that head 1-11 focuses 78% of its total attention on the amino acid *Pro* and head 12-3 focuses 27% of attention on *Phe*. Note that the maximum frequency of any single type of amino acid in the dataset is 9.4%. Detailed results for all amino acids are included in Appendix B.1.

**Figure 2:**
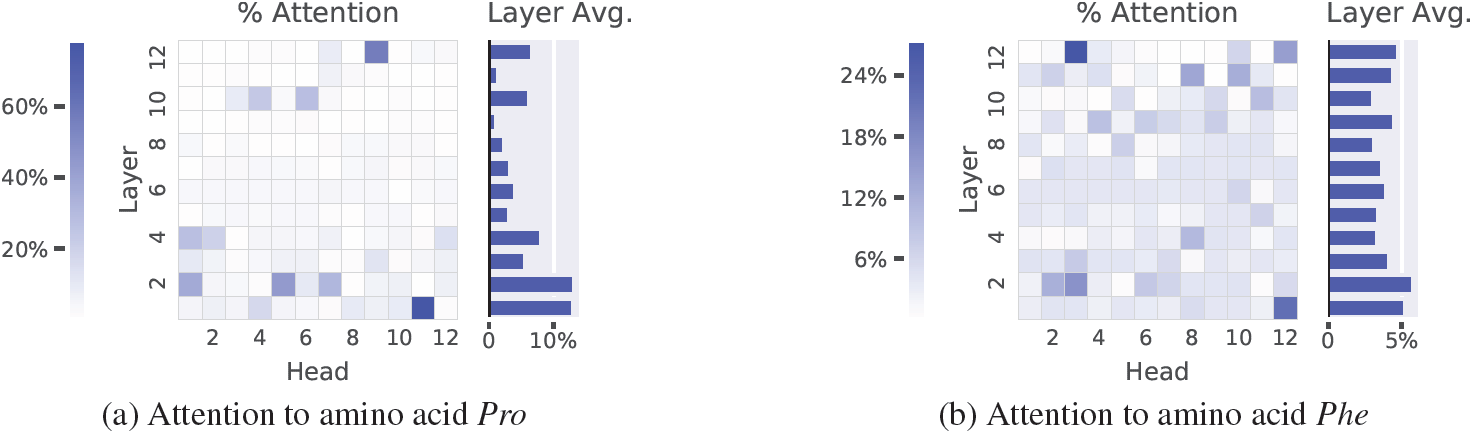
Percentage of each head’s attention that focuses on amino acids *Pro* and *Phe*, respectively, averaged over the dataset. Each heatmap cell shows the value for a single head, indexed by layer (vertical axis) and head index (horizontal axis). For example, in (a), the dark blue cell in the lower right corner shows that head 1-11 focuses 78% of its attention on *Pro*. These and other results suggest that attention heads specialize in certain types of amino acids. See Appendix B.1 for complete results.

#### Attention is consistent with substitution relationships

A natural follow-up question is whether each head has “memorized” specific amino acids to target, or whether it has actually learned meaningful properties that correlate with particular amino acids. To test the latter hypothesis, we analyze how the attention received by amino acids relates to an existing measure of structural and functional properties: the substitution matrix (see Section 2). We assess whether attention tracks similar properties by computing the similarity of *attention* between each pair of amino acids and then comparing this metric to the pairwise similarity based on the substitution matrix. To measure attention similarity, we compute the Pearson correlation between the proportion of attention that each amino acid receives across heads. For example, to measure the attention similarity between *Pro* and *Phe*, we take the Pearson correlation of the two heatmaps in Figure 2. The values of all such pairwise correlations are shown in Figure 3a. We compare these scores to the BLOSUM scores in Figure 3b, and find a Pearson correlation of 0.80, suggesting that attention is largely consistent with substitution relationships.

**Figure 3:**
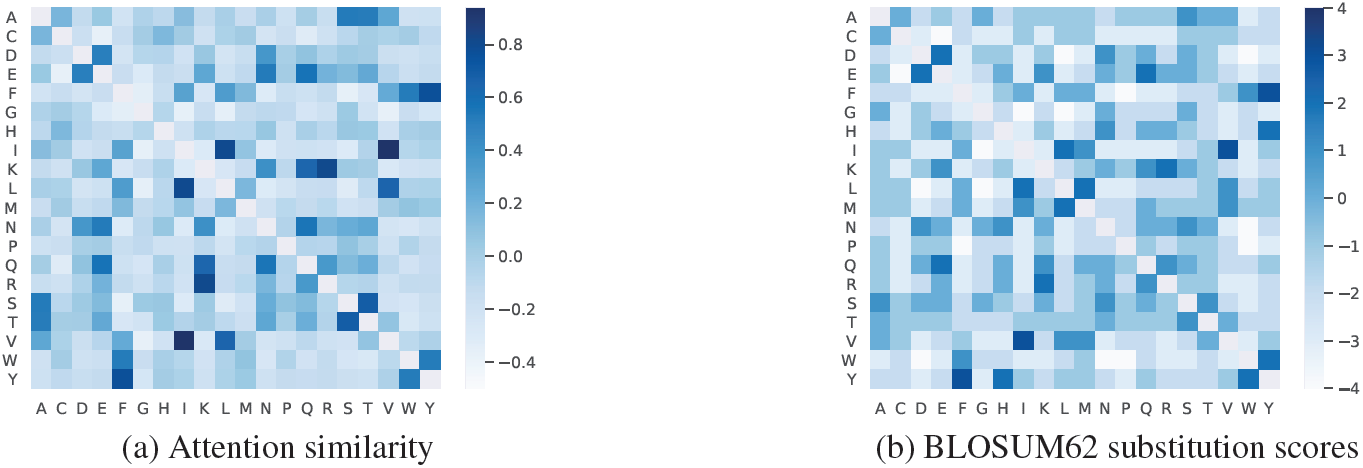
Comparison of attention similarity matrix with the substitution matrix. Each matrix entry represents an amino-acid pair (codes in Appendix B.1). The two matrices have a Pearson correlation of 0.80 with one another, suggesting that attention is largely consistent with substitution relationships.

### Protein Structure

Here we explore the relationship between attention and tertiary structure, as characterized by contact maps (see Section 2). Secondary structure results are included in Appendix B.2.

#### Attention aligns strongly with contact maps in one attention head

Figure 4 shows the percentage of each head’s attention that aligns with contact maps. A single head, 12-4, aligns much more strongly with contact maps (28% of attention) than any of the other heads (maximum 7% of attention). In cases where the attention weight in head 12-4 is greater than 0.9, the alignment increases to 76%. In contrast, the frequency of contact pairs among all token pairs in the dataset is 1.3%. Figure 1a shows an example protein and the induced attention from head 12-4.

**Figure 4:**
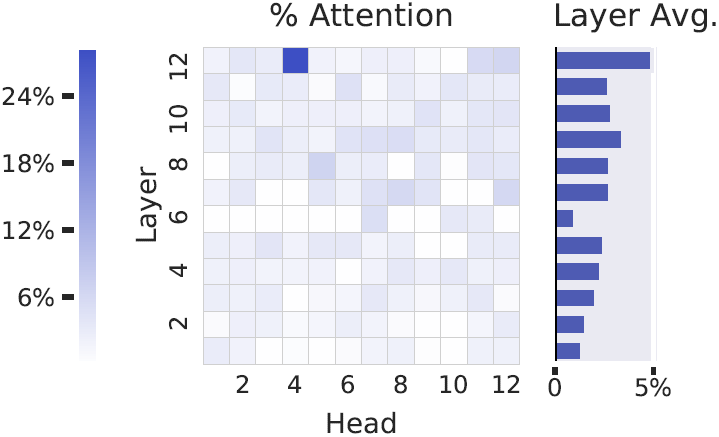
Percentage of each head’s attention that is aligned with contact maps, averaged over a dataset, suggesting that Head 12-4 is uniquely specialized for contact prediction.

Considering the model was trained with a masked language modeling objective with no spatial information in its inputs or training labels, the presence of a singular head that identifies contacts is surprising. One potential reason for this localizing behavior could be that contacts are more likely to biochemically interact with one another, thereby constraining the amino acids that may occupy these positions. In a language model, therefore, knowing contacts of masked tokens could provide valuable context for token prediction. But *how* is the model able to capture these contact relationships? One possible explanation is that the model learns the underlying geometry of the protein structure. Alternatively, the model may utilize statistical regularities resulting from the contact relationship, e.g., that amino acids in contact often coevolve. The latter explanation seems especially relevant for contacts that are far apart in the underlying sequence, since it would likely be difficult for the model to faithfully capture the geometry over a very long interval.

While there seems to be a strong correlation between the attention head output and classically-defined contacts, there are also differences. The model may have learned a differing contextualized or nuanced formulation that describes amino acid interactions. These learned interactions could then be used for further discovery and investigation or repurposed for prediction tasks similar to how principles of coevolution enabled a powerful representation for structure prediction.

#### Attention is a well-calibrated predictor of contact maps

It has been suggested that attention weights represent a model’s confidence in detecting certain features [17, 85]. We test this hypothesis with respect to contact maps by comparing attention weight in head 12-4, discussed above, with the probability of two amino acids being in contact. We estimate the contact probability by binning all amino acid pairs (*i, j*) in the dataset by their attention weights *α*_*i,j*_, and calculating the proportion of pairs in each bin that are in contact. The results are shown in Figure 5. The Pearson correlation between the estimated probabilities and the attention weights (based on the midpoint of each bin) is 0.97. This suggests that attention weight is a well-calibrated estimator in this case, providing a principled interpretation of attention as a measure of confidence.

**Figure 5:**
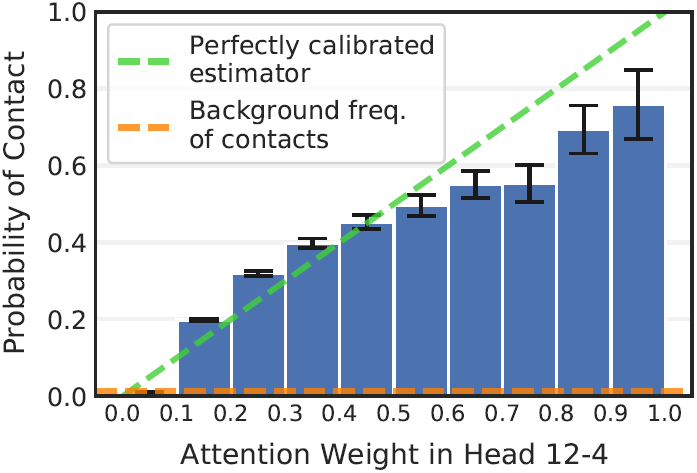
Probability two amino acids are in contact [95% confidence intervals], as a function of attention between the amino acids in Head 12-4, showing attention approximates a perfectly-calibrated estimator (green line).

### 4.3 Binding Sites

We also analyze how attention interacts with binding sites, a key functional component of proteins.

#### Attention targets binding sites, especially in the deeper layers

Figure 6 shows the proportion of attention focused on binding sites by each head. In most layers, the mean percentage across heads is significantly higher than the background frequency of binding sites (4.8%). The effect is strongest in the last 6 layers of the model, which include 15 heads that each focus over 20% of their attention on binding sites. Head 7-1, depicted in Figure 1b, focuses the most attention on binding sites (34%). Figure 7 shows the estimated probability of this head targeting a binding site, as a function of the attention weight. We also find that tokens often target binding sites from far away in the sequence. In Head 7-1, for example, the average distance spanned by attention to binding sites is 124 tokens.

**Figure 6:**
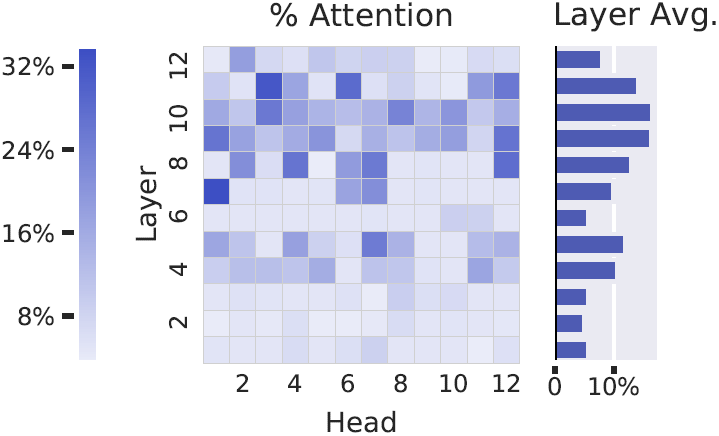
Percentage of each head’s attention that focuses on binding sites. Especially in the deeper layers, binding sites are targeted at a much higher frequency than would occur by chance (4.8%). Head 7-1 has the highest percentage (34%).

**Figure 7:**
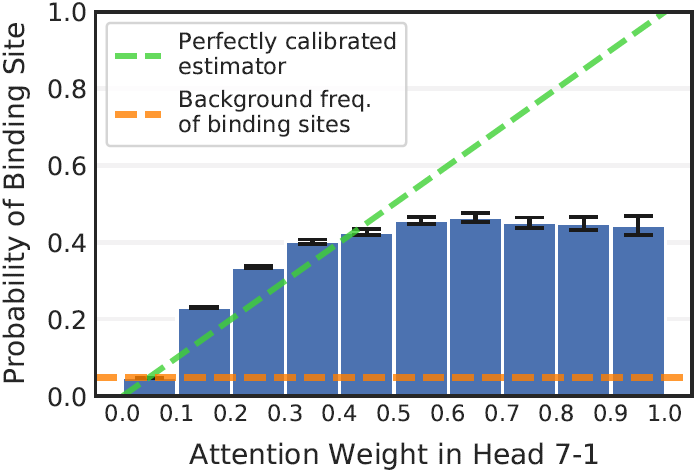
Probability that amino acid is a binding site [95% confidence intervals], as a function of attention received in Head 7-1. The green line represents a perfectly calibrated estimator.

Why does attention target binding sites, especially from long distances within the sequence? Evolutionary pressures have naturally selected proteins among the combinatorial space of possible amino acid sequences by the guiding principle that they exhibit critical function to ensure fitness. Proteins largely function to bind to other molecules, whether small molecules, proteins, or other macromolecules. Past work has shown that binding sites can reveal evolutionary relationships among proteins [42] and that particular structural motifs in binding sites are mainly restricted to specific families or superfamilies of proteins [37]. Thus binding sites provide a high-level characterization of the protein that may be relevant for the model throughout the sequence.

### 4.4 Cross-Layer Analysis

Here we analyze how attention captures properties of varying complexity across different layers of the model, and we compare these results to a probing analysis of layer outputs (see Section 3).

#### Attention targets higher-level properties in deeper layers

As shown in Figure 8, deeper layers focus relatively more attention on binding sites and contacts (high-level concept), whereas secondary structure (low-to mid-level concept) is targeted more evenly across layers. The probing analysis (Figure 9) similarly shows that the model first forms representations of secondary structure before fully encoding contact maps and binding sites. This result is consistent with the intuition that the model must understand local structure before it can form representations of higher-order structure and function. Prior work in NLP also suggests that deeper layers in text-based Transformers attend to more complex properties [83] and encode higher-level representations [36, 56, 58, 76].

**Figure 8:**
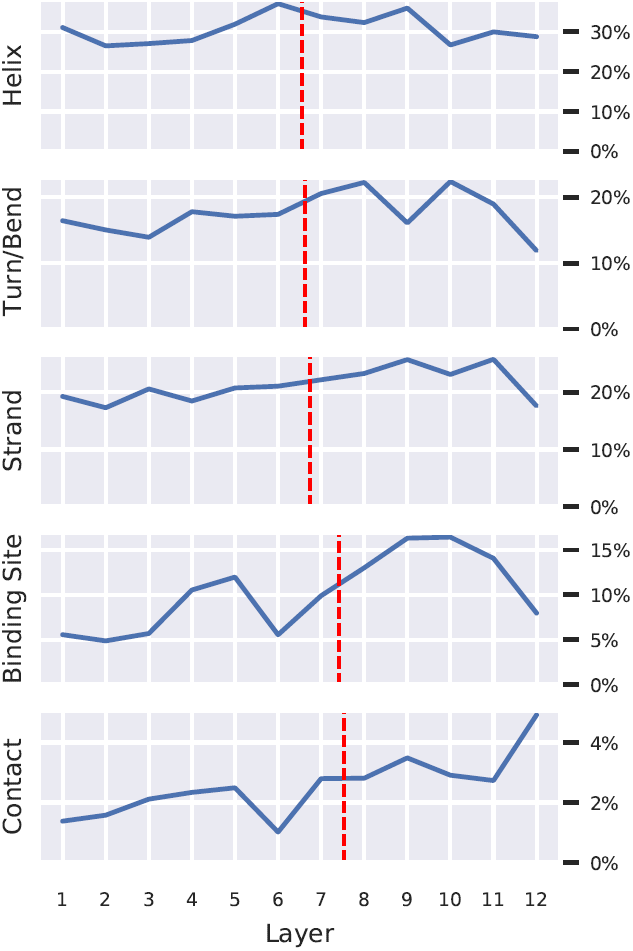
Each plot shows the percentage of attention focused on the given property, averaged over all heads within each layer. The plots, sorted by center of gravity (red dashed line), show that heads in deeper layers focus relatively more attention on binding sites and contacts, whereas attention toward specific secondary structures (*Strand, Turn/Bend, Helix*) is more even across layers. More granular analyses of these properties can be found in Figures 4 and 6 and Appendix B.2.

**Figure 9:**
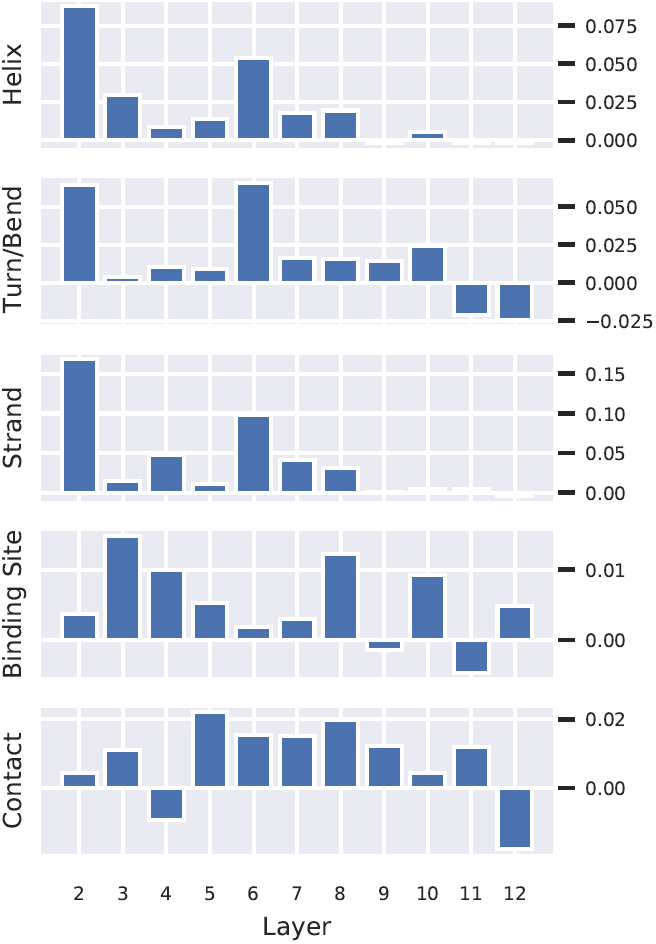
Differential performance of diagnostic classifier by layer, sorted by task order in Figure 8. Each plot shows the change in performance between the given layer and the previous layer. For secondary structure (*Strand, Turn/Bend, Helix*) the increases occur primarily in the first 5 layers, while increases for binding sites and contact maps are spread more evenly across layers. See Section 3 for a description of evaluation metrics. Absolute performance for each task included in Appendix C.

However, one seemingly contrary result is the drop in performance of the contact map classifier in the final layer (Figure 9). This may be explained by the fact that the final layer outputs are not used to compute attention, but rather as an input to the final classifier layer that predicts the masked token for the language model. Whereas attention is a property of token pairs, the predictions are specific to an individual token. Therefore, contact relationships, which exist between tokens, may not be as useful to the model in the final layer. Practitioners may take note to consider the second to last layer when generating embeddings for downstream pairwise tasks.

## 5. Related Work

Here we contextualize our techniques and findings in the protein modeling and interpretable neural network literatures, especially in natural language processing.

### 5.1 Protein language models

There is a vast literature dedicated to advancing machine learning (ML) algorithms for proteins. More specifically, deep neural networks for protein language modeling have received broad interest. Early work in this direction applied the Skip-gram model [50] to construct continuous embeddings from protein sequences for downstream tasks [7]. Sequence-only language models have since been trained through autoregressive or autoencoding self-supervision objectives for both discriminative and generative tasks, for example, using LSTMs or Transformer-based architectures [2, 9, 60, 64].[64] showed that the output embeddings from a pretrained Transformer model could be transformed to predict structural and functional properties of proteins, but attention weights were not explored in this analysis. TAPE created a benchmark of five tasks ranging from remote homology to fluorescence prediction to assess protein representation learning models. In [48, 63], autoregressive generative models were trained to predict the functional effect of mutations and generate natural-like proteins. All aforementioned studies have been the subject of rapid development, with Transformer architectures seemingly providing the most promising avenue for future research. In addition to sequence-only approaches, Transformer architectures have been adapted to incorporate structural information [34].

### 5.2 Interpreting Models in NLP

The rise of deep neural networks in ML has also led to much work on interpreting these so-called black-box models. Interpretability is a broad topic in ML with many facets [20, 46]; our present work focuses on post hoc interpretation. This section reviews the NLP interpretability literature, which is directly comparable to our work on interpreting protein language models.

#### General interpretability

Interpretability techniques in NLP mainly focus on explaining instance-level predictions. Influence functions explain a model’s prediction on a given instance by identifying the influential training examples for that prediction [14, 39]. Methods that learn to explain by extracting rationales from the input and only using those rationales to make a prediction are *faithful* by design [8, 13, 43, 62]. Model-agnostic methods that are not based on gradients do not require model differentiability. Examples include approaches that fit a linear classifier to a model’s prediction on perturbations of the input [4, 61, 72]. Yet another line of research deploys post hoc rationalization for explaining model behavior. Gradient-based techniques are the most commonly used methods in this category [25, 73, 74]. Other approaches include using language models for generating explanations [53, 59].

#### Interpreting Transformers

The Transformer is a neural architecture that uses attention to accelerate learning [80]. Transformers are the backbone of state-of-the-art pre-trained language models in NLP including BERT [19]. BERTology focuses on interpreting what the BERT model learns about natural language by using a suite of probes and interventions [65]. So-called *diagnostic classifiers* are used to interpret the outputs from BERT’s layers [81].

At a high level, mechanisms for interpreting BERT can be categorized into three main categories: interpreting the learned embeddings [1, 16, 23, 49, 86], BERT’s learned knowledge of syntax [27, 30,32, 45, 47, 76], and BERT’s learned knowledge of semantics [24, 76].

#### Interpreting attention specifically

In this paper, we focused on the interpretability of Transformer language models as they apply to protein sequences, with a specific focus on the attention mechanism. Interpreting attention on natural language sequences is a well-established area of research [12, 30, 87, 90]. In some cases, it has been shown that attention correlates with syntactic and semantic relationships in natural language [15, 32, 83]. To our knowledge, no previous work has examined whether attention is a *well-calibrated* estimator of these properties, though previous work as explored the calibration of Transformer outputs from an interpretability perspective [18].

Depending on the task and model architecture, attention may have more or less explanatory power for model predictions [35, 51, 57, 71, 79]. Visualization techniques have been used to convey the structure and properties of attention in Transformers [31, 40, 80, 82]. Recent work has begun to apply attention to guide mapping of sequence models outside of the domain of natural language [70].

Research on using pre-trained language models for protein sequences has been motivated by drawing parallels with natural language. Our work also draws motivation from interpretability techniques in NLP to uncover the properties captured by protein language models. The availability of TAPE suite of protein classification tasks inspired us to use diagnostic classifiers for probing of BERT’s layers.

## 6 Conclusions and Future Work

This paper builds on the synergy between NLP and computational biology by adapting and extending NLP interpretability methods to protein sequence modeling. We show how a Transformer language model recovers structural and functional properties of proteins and integrates this knowledge directly into its attention mechanism. Our analysis reveals that attention not only captures properties of individual amino acids, but also discerns more global properties such as binding sites and tertiary structure. In some cases, attention also provides a well-calibrated measure of confidence in the model’s predictions of these properties. We hope that machine learning practitioners can apply these insights in designing the next generation of protein sequence models.

The present analysis identifies associations between attention and various properties of proteins. It does not attempt to establish a causal link between attention and model behavior [28, 84], nor to *explain* model predictions [35, 87]. While the focus of this paper is reconciling attention patterns with known properties of proteins, one could also leverage attention to discover novel types of properties and processes. In this way, unsupervised methods like language modeling can serve as tools for scientific discovery. For example, one may be able to discover new insights into evolutionary history by better understanding phylogenetic relationships [89] or into development by understanding ontogenic relationships among cells (recently studied using single-cell RNA sequence data [88] rather than proteins). But in order for learned representations to be useful to domain experts, they must be presented in an appropriate context to facilitate discovery. Visualizing attention in the context of protein structure (Figure 1) is one example of this approach. We believe there is potential to develop other such contextual visualizations of learned representations in biology and other domains.

## 7 Broader Impact

Interpretability is the hallmark of trustworthy artificial intelligence and is especially important when technologies are directly used by people in consequential activities such as protein engineering or scientific discovery. This work illuminates some of the inner workings of large-scale language models for proteins, which are otherwise inscrutable, and may therefore facilitate their use by scientists and engineers. Many of the safety and governance considerations for pretrained language models for human language also apply for protein language [78], e.g. disparate resources available for training, fine-tuning, and using these models. The governance recommendations of post-market surveillance and co-responsibility among producers and users may be applied to mitigate these potential issues.

Scientists and engineers may pursue beneficent applications of protein language models such as the design of novel disease therapies, antibodies for novel infectious diseases, or food ingredients for low-allergenic infant formulas. They may also, however, pursue hostile applications such as the design of protein toxins [77], which, in the case of developing biological weapons, could have significant geopolitical ramifications. Various strategies could be applied to mitigate such risks, including broad-based governmental regulation, shared responsibility between users and producers of technology, and post-market surveillance.

Since the molecular biology revolution in the 1960s, there has been a large focus on a small number of model organisms, such as *C. elegans, Drosophila melanogaster*, and *Mus musculus*, as part of biological research [5, 44]. The protein databases that are used to train protein language models may therefore have unknown biases across the tree of life. These dataset biases may be reflected in the learned models, and further in their interpretation.

## 8. Acknowledgements

We would like to thank Xi Victoria Lin, Stephan Zheng, and Melvin Gruesbeck for their valuable feedback.

### A Modeling and Experimental Details

#### A.1 BERT Transformer Architecture

##### Stacked Encoder

BERT uses a stacked-encoder architecture, which inputs a sequence of tokens ***x*** = (*x*_*1*_, …, *x*_*L*_) and applies position and token embeddings followed by a series of encoder layers. Each layer applies multi-head self-attention (see below) in combination with a feedforward network, layer normalization, and residual connections. The output is a sequence of continuous embeddings ***z*** = (*z*_*1*_, …, *z*_*L*_).

##### Self-Attention

Given an input ***x*** = (*x*_*1*_, …, *x*_*L*_), the self-attention mechanism assigns to each token pair *i, j* an attention weight *α*_*i,j*_ *>* 0 where ∑ _*j*_ *α*_*ij*_ = 1. Attention in BERT is bidirectional. In the multi-layer, multi-head setting, *α* is specific to a layer and head. The BERT-Base model has 12 layers and 12 heads.

The attention weights *α*_*i,j*_ are computed from the scaled dot-product of the *query vector* of *i* and the *key vector* of *j*, followed by a softmax operation. The attention weights are then used to produce a weighted sum of value vectors:

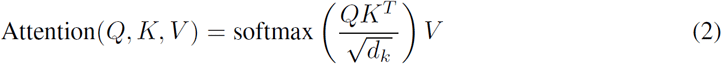

using query matrix *Q*, key matrix *K*, and value matrix *V*, where *d*_*k*_ is the dimension of *K*. In a multi-head setting, the queries, keys, and values are linearly projected *h* times, and the attention operation is performed in parallel for each representation, with the results concatenated.

#### A.2 Datasets

We used two protein sequence datasets from the TAPE repository for the analysis: the ProteinNet dataset [3, 10, 26, 52] and the Secondary Structure dataset [10, 38, 52, 60]. The former was used for analysis of amino acids and contact maps, and the latter was used for analysis of secondary structure. We additionally created a third dataset for binding site analysis from the Secondary Structure dataset, which was augmented with binding site annotations obtained from the Protein Data Bank’s Web API.^3^ We excluded any sequences for which binding site annotations were not available. The resulting dataset sizes are shown in Table 10. For the analysis of attention, a random subset of 5000 sequences from the training split of each dataset was used, as the analysis was purely evaluative. For training and evaluating the diagnostic classifier, the full training and validation splits were used.

### B. Additional Results of Attention Analysis

#### B.1 Amino Acids

Figures 11-13 show attention analysis results for all amino acids. Amino acid codes are found in Figure 14.

**Figure 10:**
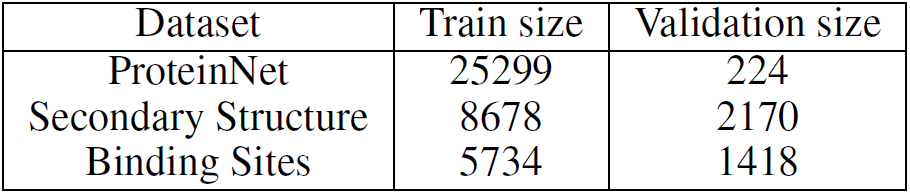
Datasets used in analysis

**Figure 11:**
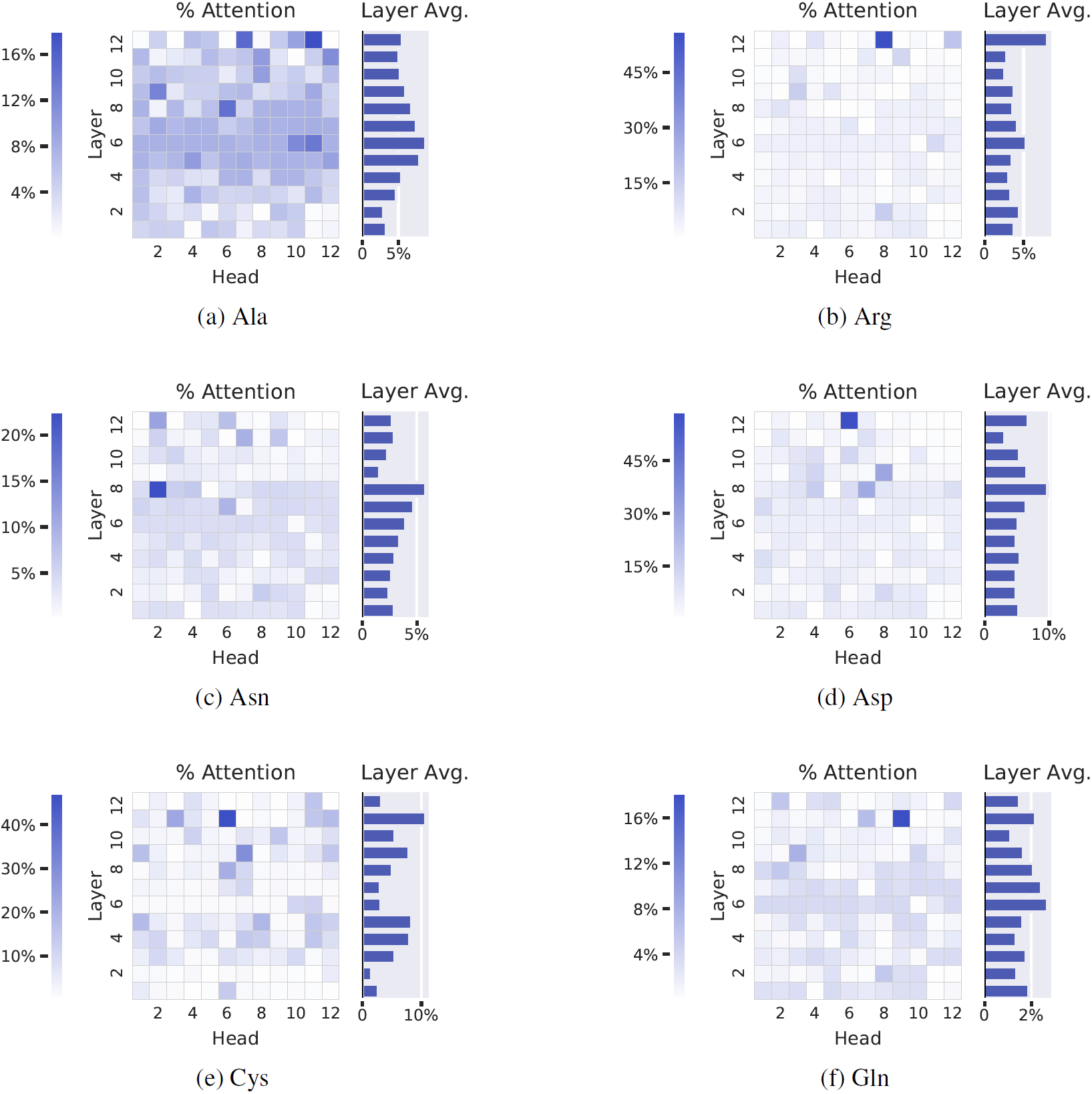
Percentage of each head’s attention that is focused on the given amino acid, averaged over a dataset. Each entry in the heatmap shows the value for a single head, while the bar plot on the right of each figure shows the layer average across heads.

**Figure 12:**
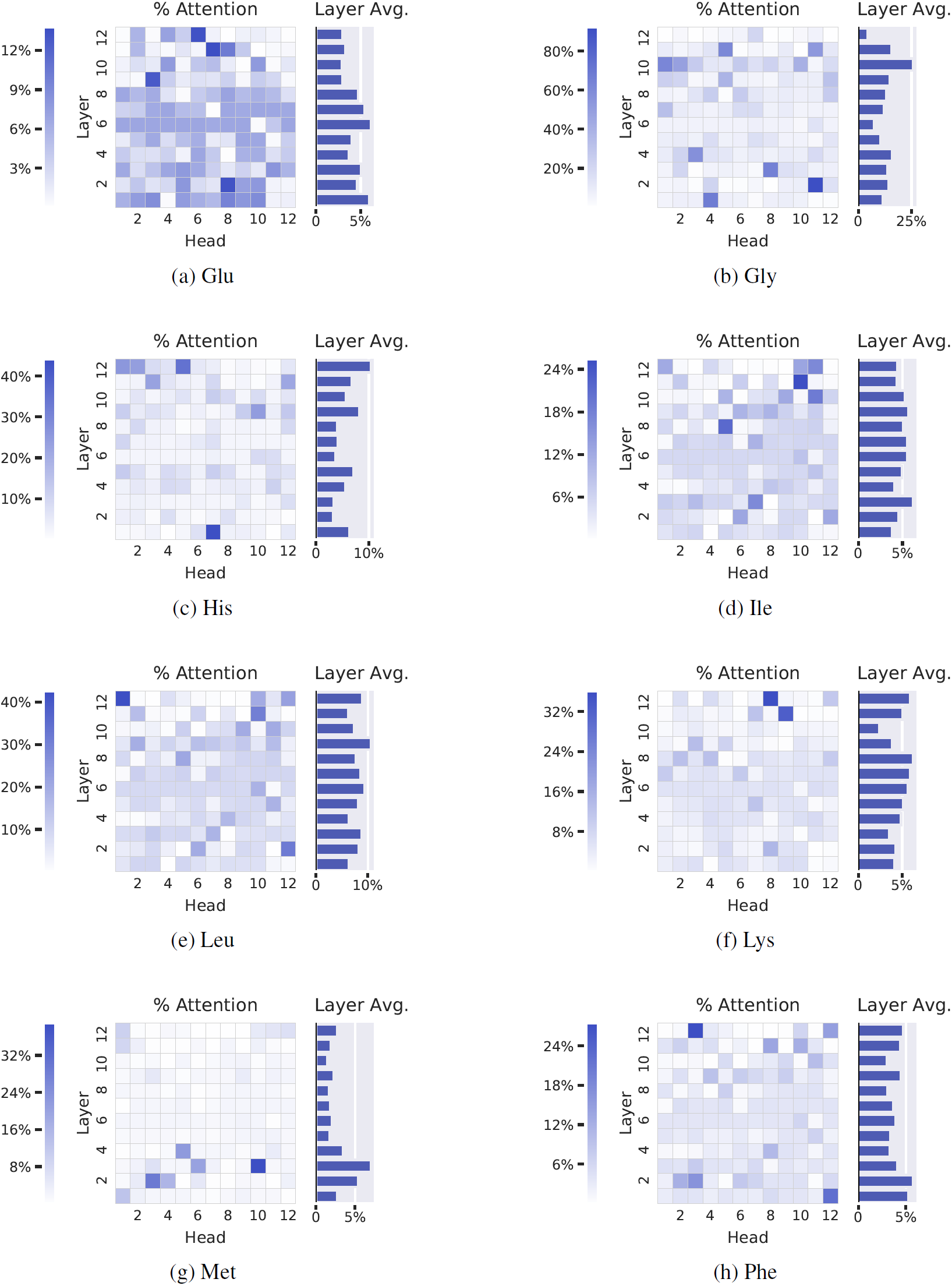
Percentage of each head’s attention that is focused on the given amino acid, averaged over a dataset. Each entry in the heatmap shows the value for a single head, while the bar plot on the right of each figure shows the layer average across heads. (cont.)

**Figure 13:**
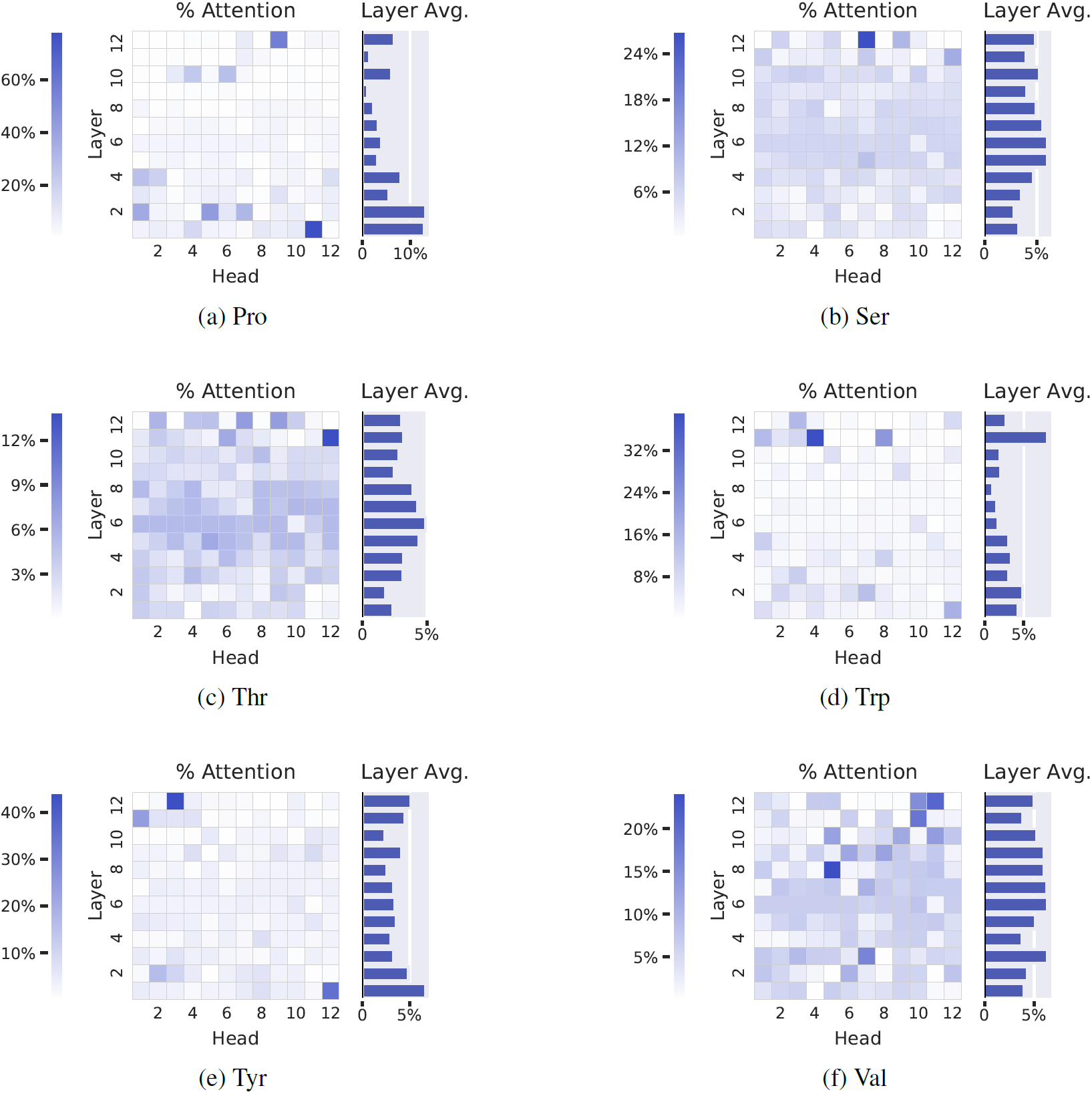
Percentage of each head’s attention that is focused on the given amino acid, averaged over a dataset. Each entry in the heatmap shows the value for a single head, while the bar plot on the right of each figure shows the layer average across heads. (cont.)

**Figure 14:**
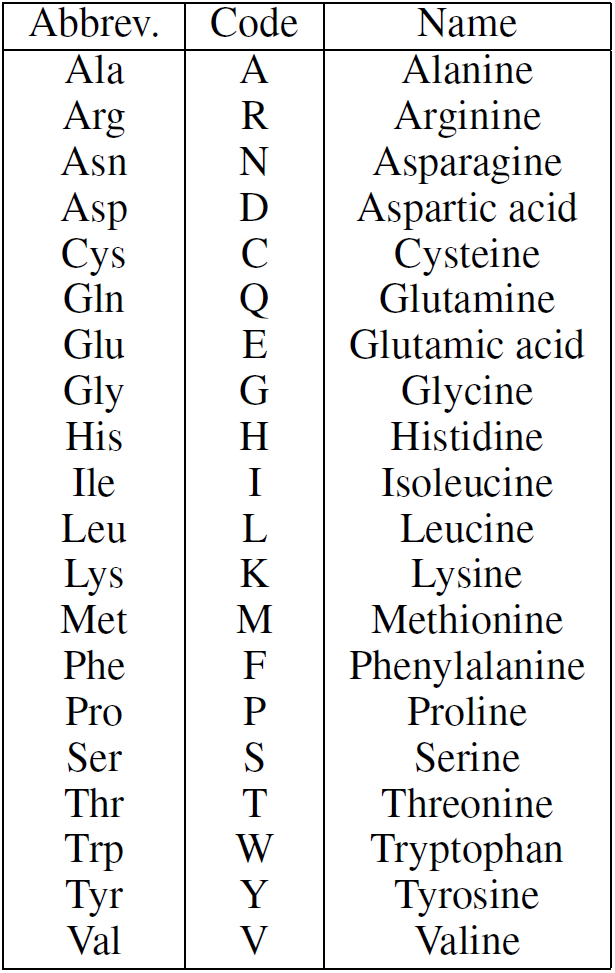
Amino acids and associated codes.

#### B.2 Secondary Structure

**Figure 15:**
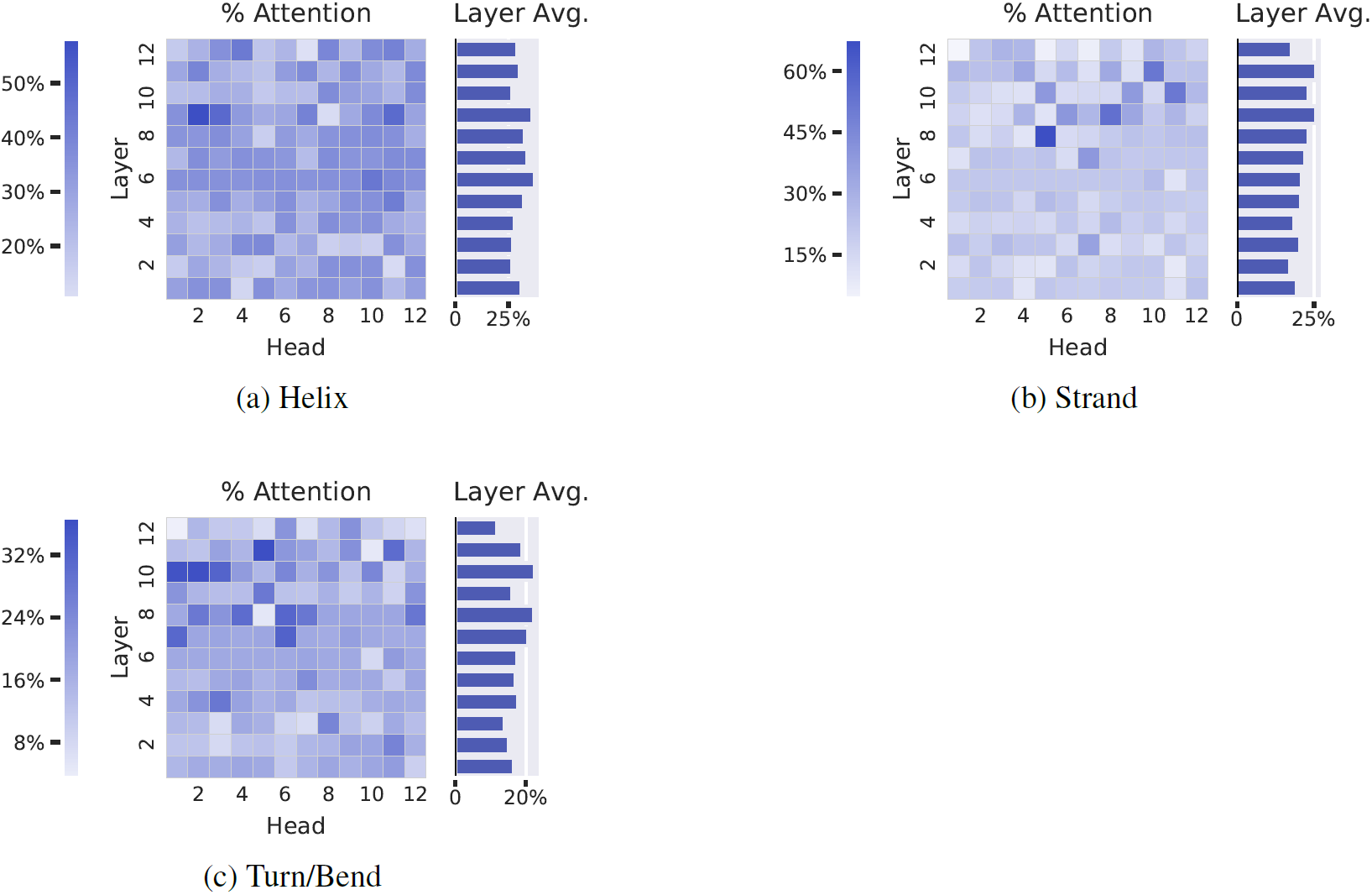
Percentage of each head’s attention that is focused on the given secondary structure type, averaged over a dataset. Each entry in the heatmap shows the value for a single head, while the bar plot on the right of each figure shows the layer average across heads.

#### C Detailed Probing Results

Preprint. Under review.

**Figure 16:**
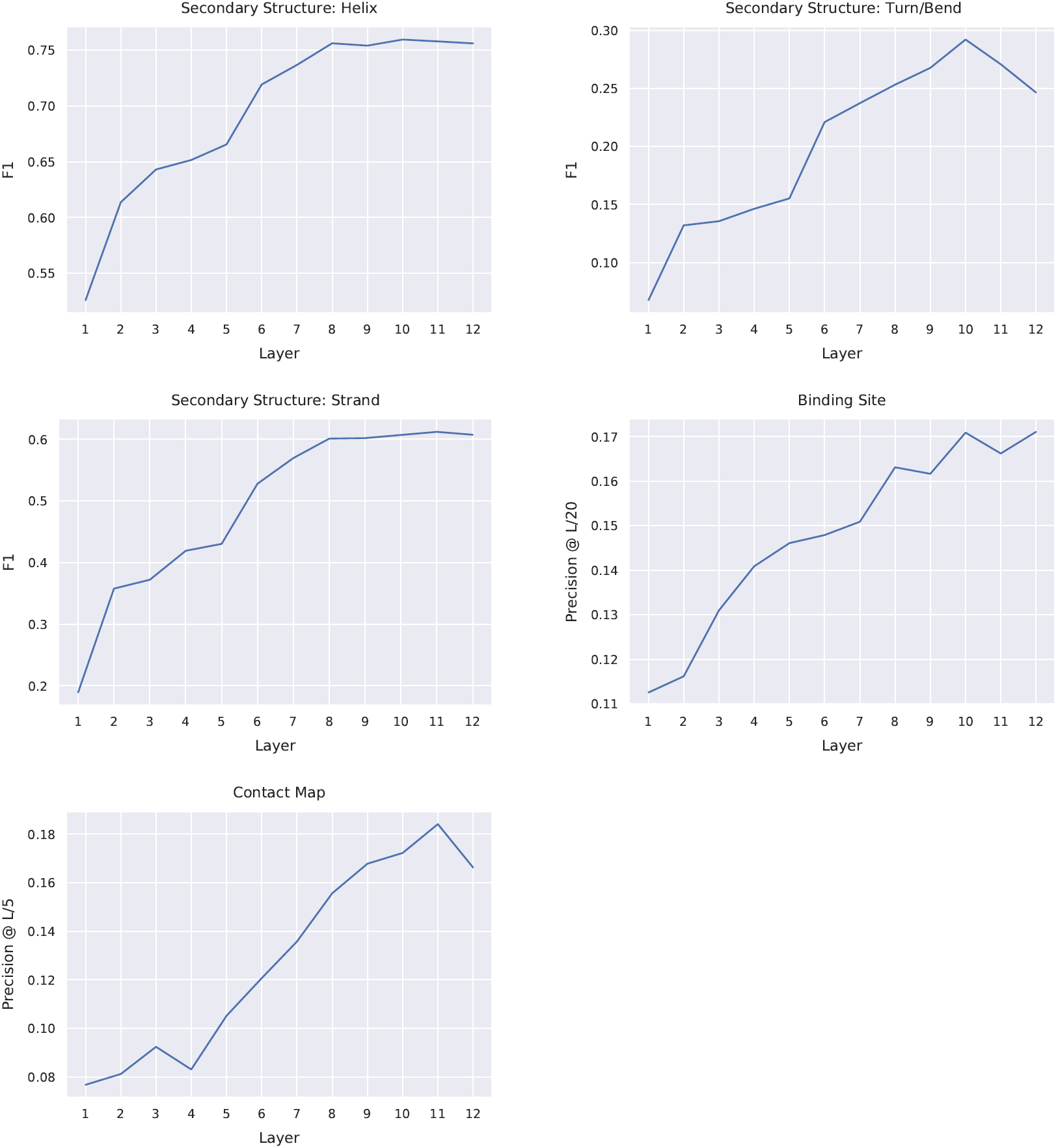
Diagnostic classifier performance for tasks described in Figure 9. For secondary structure prediction, we measure F1 score; for contact map prediction; we measure precision@*L/*5, where *L* is the length of the protein sequence, following standard practice [52]; for binding site prediction, we measure precision@*L/*20, since approximately one in twenty amino acids in each sequence is a binding site (4.8%). The raw scores of the different metrics cannot be compared directly; the goal of this analysis is to compare the relative shape of the performance curves across tasks.

## Footnotes

1 https://github.com/songlab-cal/tape

2 94% of sequences had fewer than 512 amino acids.

3 http://www.rcsb.org/pdb/software/rest.do

